# The new silicone N99 half-piece respirator, VJR-NMU N99: A novel and effective tool to prevent COVID-19

**DOI:** 10.1101/2020.07.23.217372

**Authors:** Anan Manomaipiboon, Sujaree Pupipatpab, Pongsathorn Chomdee, Pathiporn Boonyapatkul, Thananda Trakarnvanich

## Abstract

Filter facepiece respirators (FFRs) are critical for preventing the transmission of respiratory tract infection disease, especially the dreadful coronavirus 2 (SARs-CoV-2). The N95 mask is a prototype, high-efficiency protective device that can effectively protect against airborne pathogens of less than 0.3 μm. The N95 mask is tightly fitting and has high filtration capacity. The ongoing COVID-19 pandemic has led to a greater requirement for FFR. This rising demand greatly exceeds current production capabilities and stockpiles, resulting in shortages. To address this, our team has invented a new type of half-piece respirator made from silicone and assembled with HEPA or elastostatic filter. A variety of methods have been used to evaluate this new device, including a qualitative fit test with the Bitrex® test kit and filtration test. The preliminary results showed that the new N99 respirators pass the fit test. The filtration tests also confirmed the superiority of N99 over traditional N95 masks, with a mean performance of protection greater than 95%. For the filters, we used two types: SafeStar, which is a kind of HEPA filter; and CareStar, which is considered an elastostatic filler. CareStar was developed to filter virus and bacteria in the operating room, with a limit duration of use up to 24 h, while the safe star was designed for 72 h use and has the quality equivalent to a HEPA filter. Our study demonstrated superior filtration efficacy of both filters, more than 98% even after 24 h of use. CareStar has significantly more filtration efficacy than a safe star (p < 0.001). In conclusion, the development of our new N99 half-piece respirator should ultimately be applicable to healthcare workers with at least non-inferiority to the previously used N 95 respirators.. Currently, the adequate supply of such equipment is not feasible. The advent of the new protective device will help protect healthcare workers and replenish the shortage of N95 respirators during the COVID-19 pandemic.

## Introduction

Since the rapid spread of SARs-CoV-2 worldwide, resulting in the novel coronavirus disease 2019 (COVID-19) pandemic, the shortage of personal protective equipment (PPE), including surgical masks and N95 respirators, has been a serious concern [1]. The transmission of SARs-CoV-2 can occur by contact or droplets released from infected persons when coughing, talking, and sneezing [2]. Airborne transmission might occur during aerosol-generating procedures, such as tracheal intubation [3]. Therefore, health care personnel (HCP), especially frontline workers, should be properly protected with appropriate equipment. A filtering facepiece and N95 respirators are used as high-performance filtering masks to protect staff against both droplets and aerosols [4]. To avoid cross contamination, these devices are designed to be disposable after a single use [5]. Consequently, the consumption of FFP masks has been overwhelmed, and a supply shortage for HCP has already been reported in many countries [6].

To meet the need for FFRs during a pandemic, Navamindradhiraj University has invented a new model of silicone N99 facepiece respirator by using the silicone mask and the HEPA filter normally used in the operating room (CareStar or SafeStar, Draeger, Germany) (Fig 1). This filter has a minimum filtration efficiency of ≥ 99% to the challenge aerosols consisting of particles in the most penetrating particle size range (approximately 0.3 μm). This can prevent penetration of viruses and bacteria and improves on the protective efficacy of the traditionally used N95 respirators. Here we describe a novel silicone N95 respirator aimed at HCPs who are in direct contact with patients, and capable of contributing to the replenishment of N95 masks.

**Fig 1.**
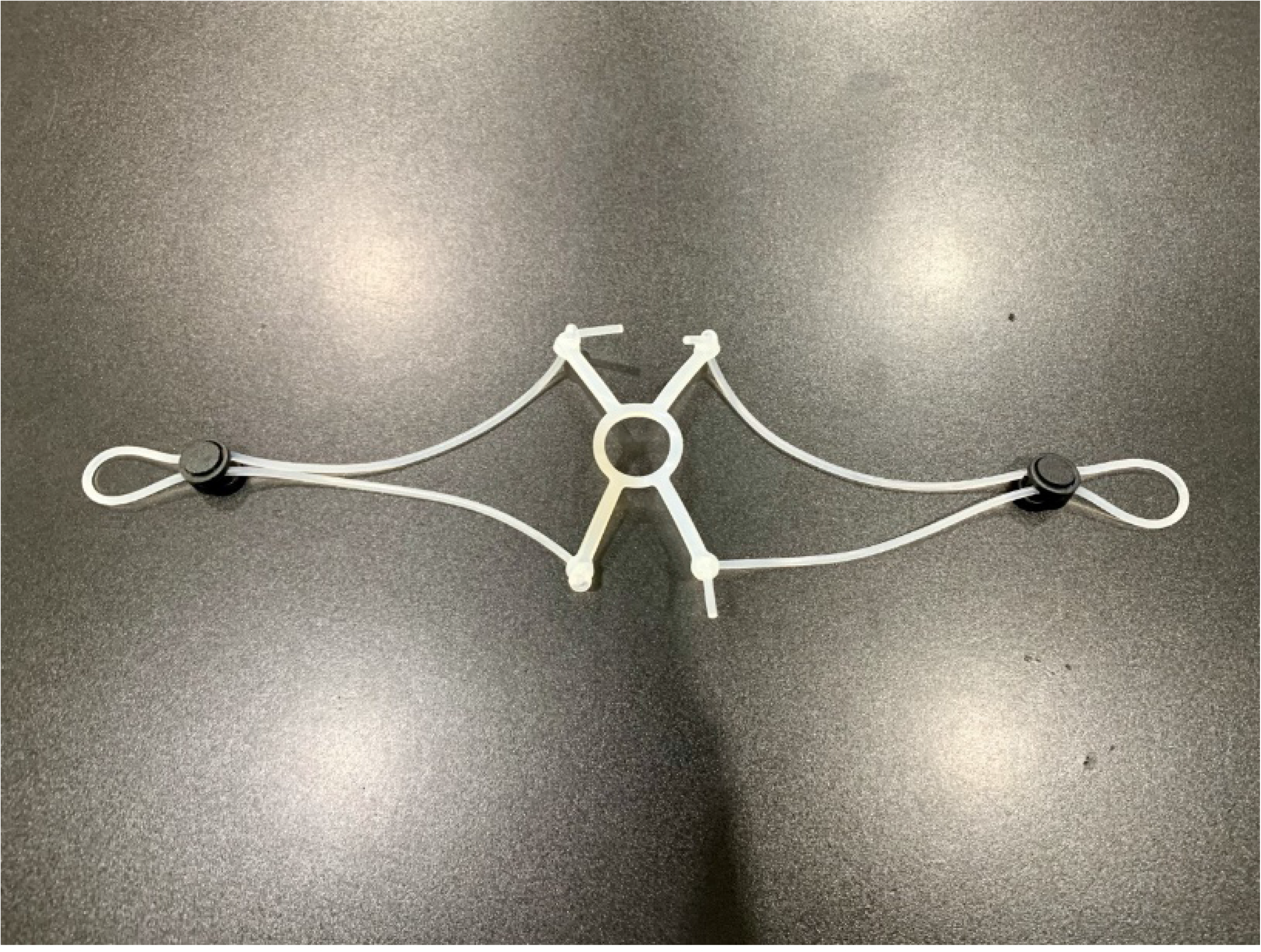
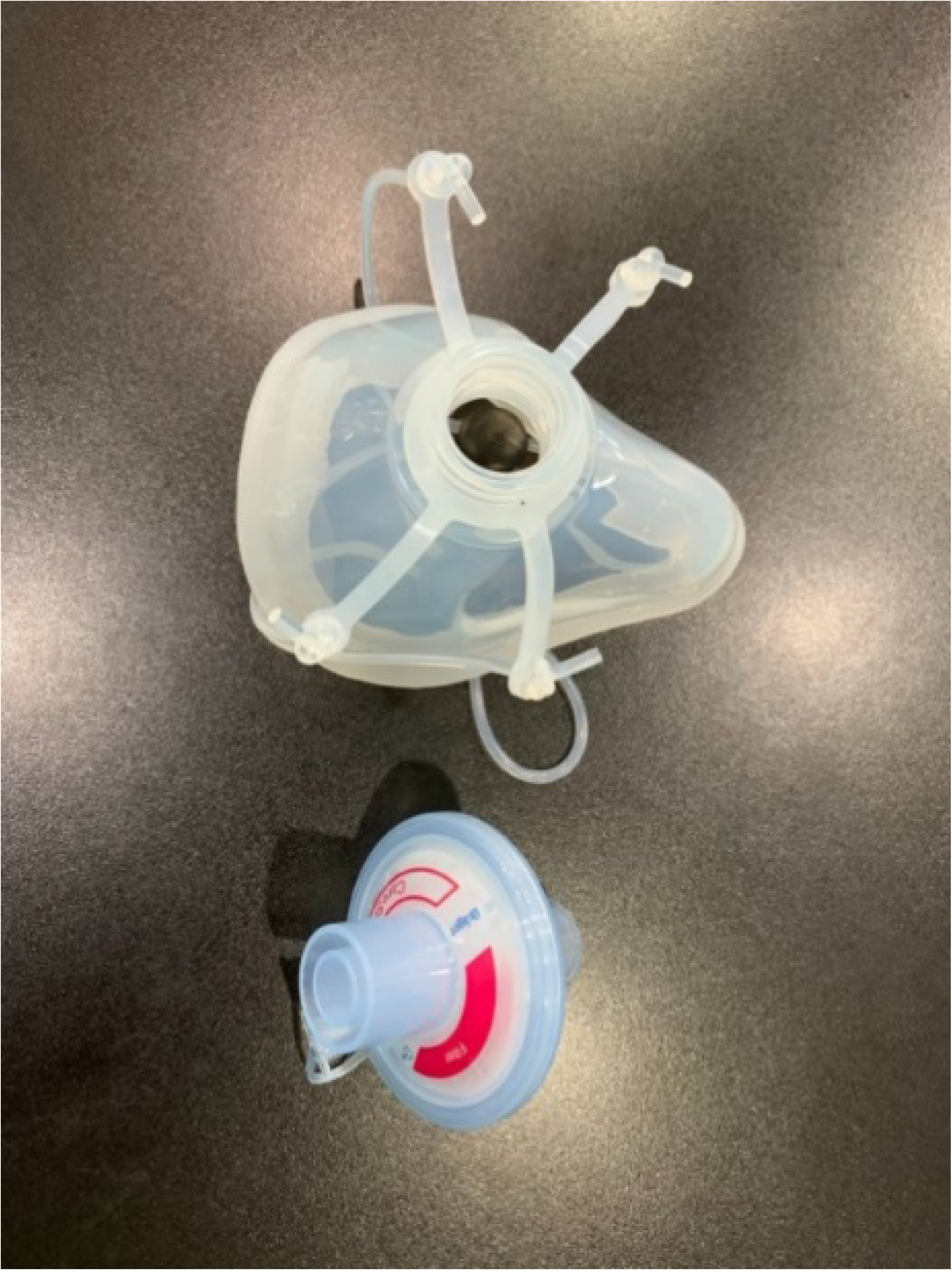
Silicone N99 Mask.

An important part of the silicone mask is the O-ring strap which can be adjusted to prevent face-seal leakage (Fig 2).

**Fig 2.**
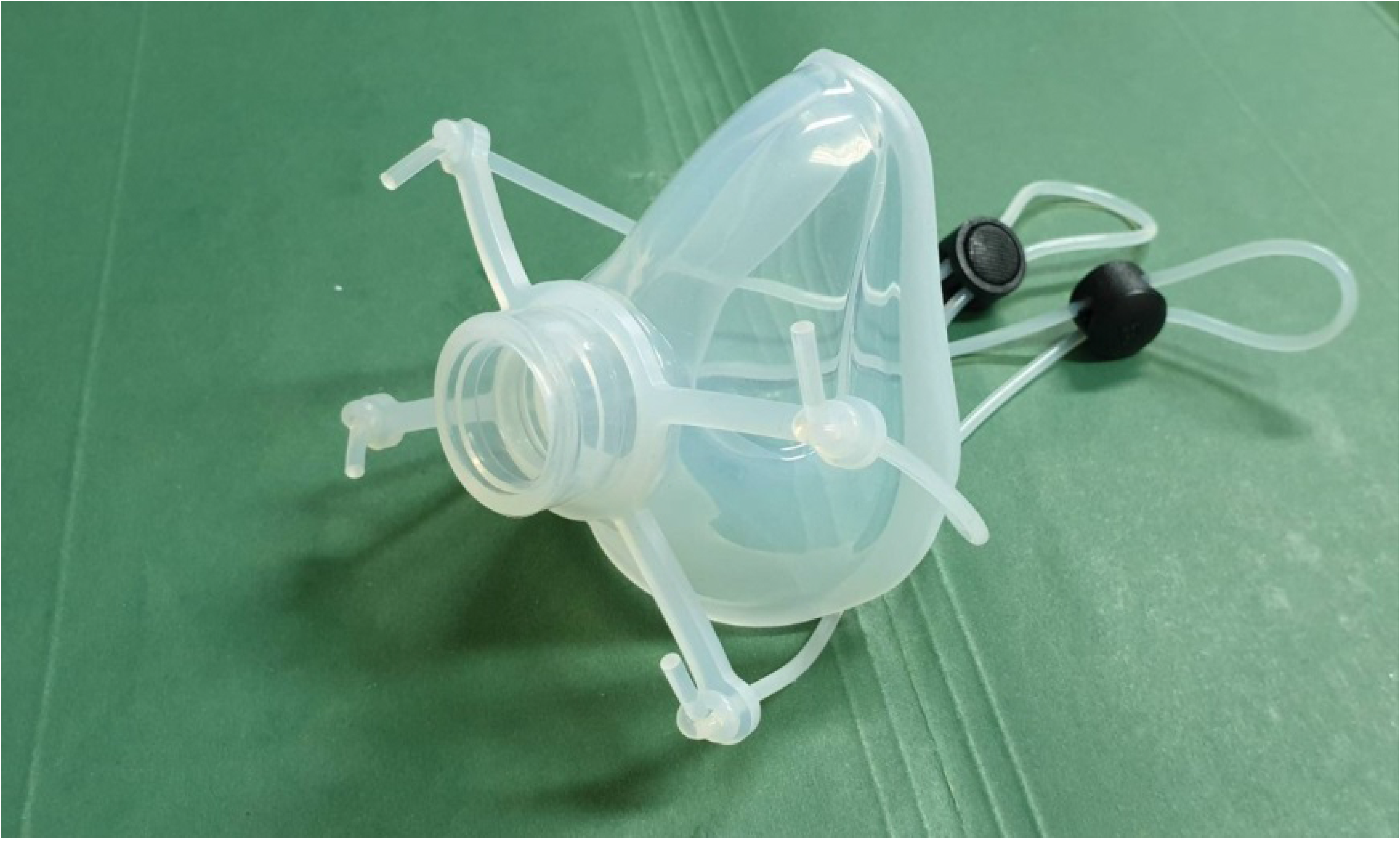
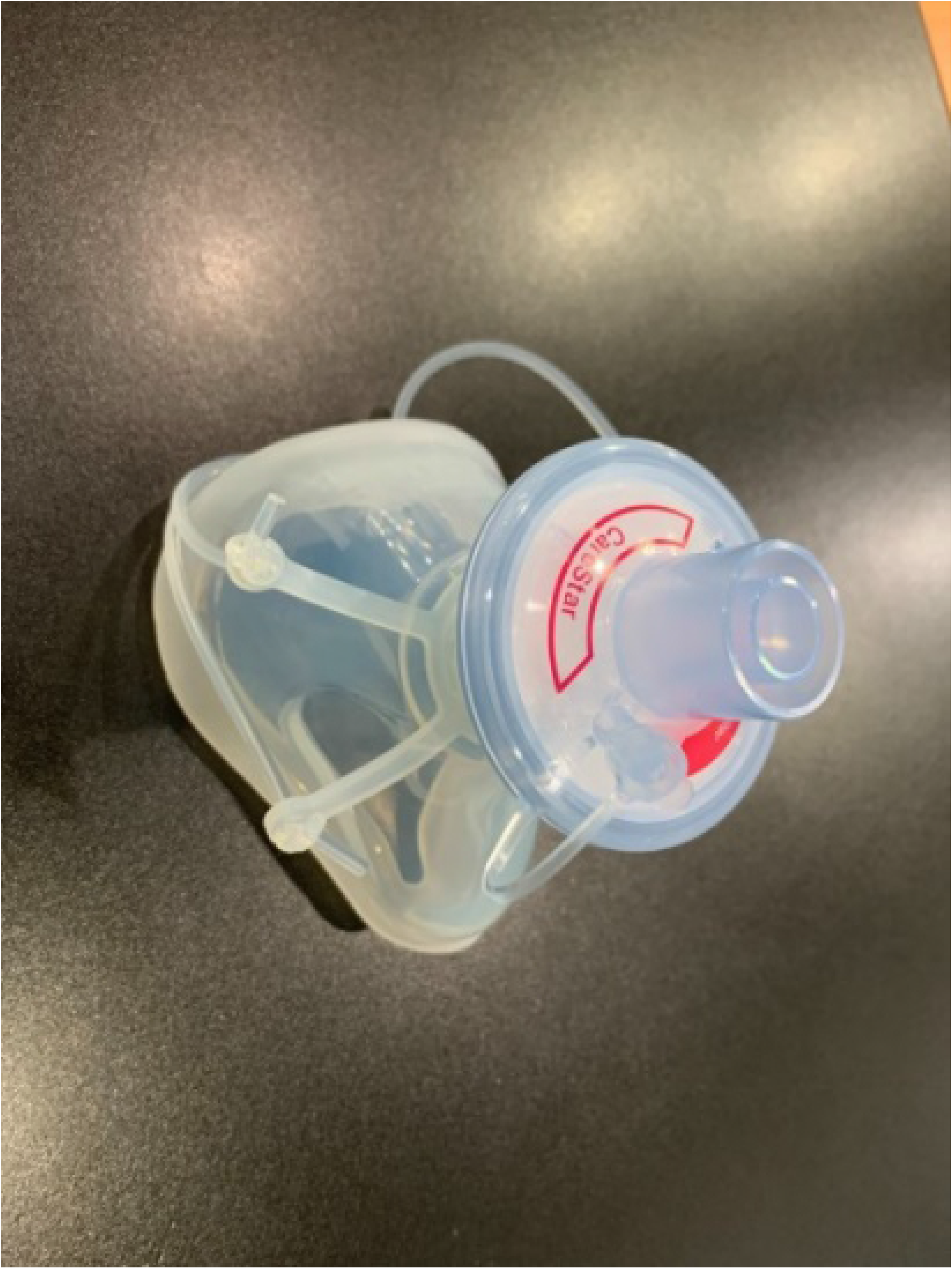
Components of the Silicone N99. The respirator consists of a silicone mask and filter.

## Components of the N99 mask

The silicone N99 mask is made using Silibione MM Series 71791 U silicone (Bluestar silicones, Shanghai, China), which are elastomers comprising of polymethyl vinyl siloxane gums and silica. In this particular series, this silicone rubbers are cured after the addition of a vulcanizing agent (chosen as a function of the production process). The heat cure is done after the addition of an organic peroxide compound and post-curing at 200°C after vulcanization. This series includes four products that differ by their hardness once cured: 40, 50, 60, and 70 types (data in the supplemental file). Once processed, the Silibione MM series 71791 U are intended for food contact and biomedical applications. The advantages of the Silibione MM series 71791 U include easy processing, highly transparent, and excellent mechanical properties (including high tear strength and a good compromise between tear strength and compression set). The Silibione MM series 71791 U is also very resistant to oxidizing agents for all sterilization modes, and is chemically inert. Moreover, samples of the silicone MM series have been subjected to migration heats in accordance with European and American regulations. The masks are made in three sizes: small, medium, and large. We assembled the silicone with a HEPA filter (CareStar® electrostatic filter), which is normally used together with a ventilator to filter bacteria and viruses from airborne transmission (Fig 2). The efficiency of the silicone mask is based on the efficiency of the air filter, which is considered a high-performing electrostatic filtration medium (Fig 3).

**Fig 3.**
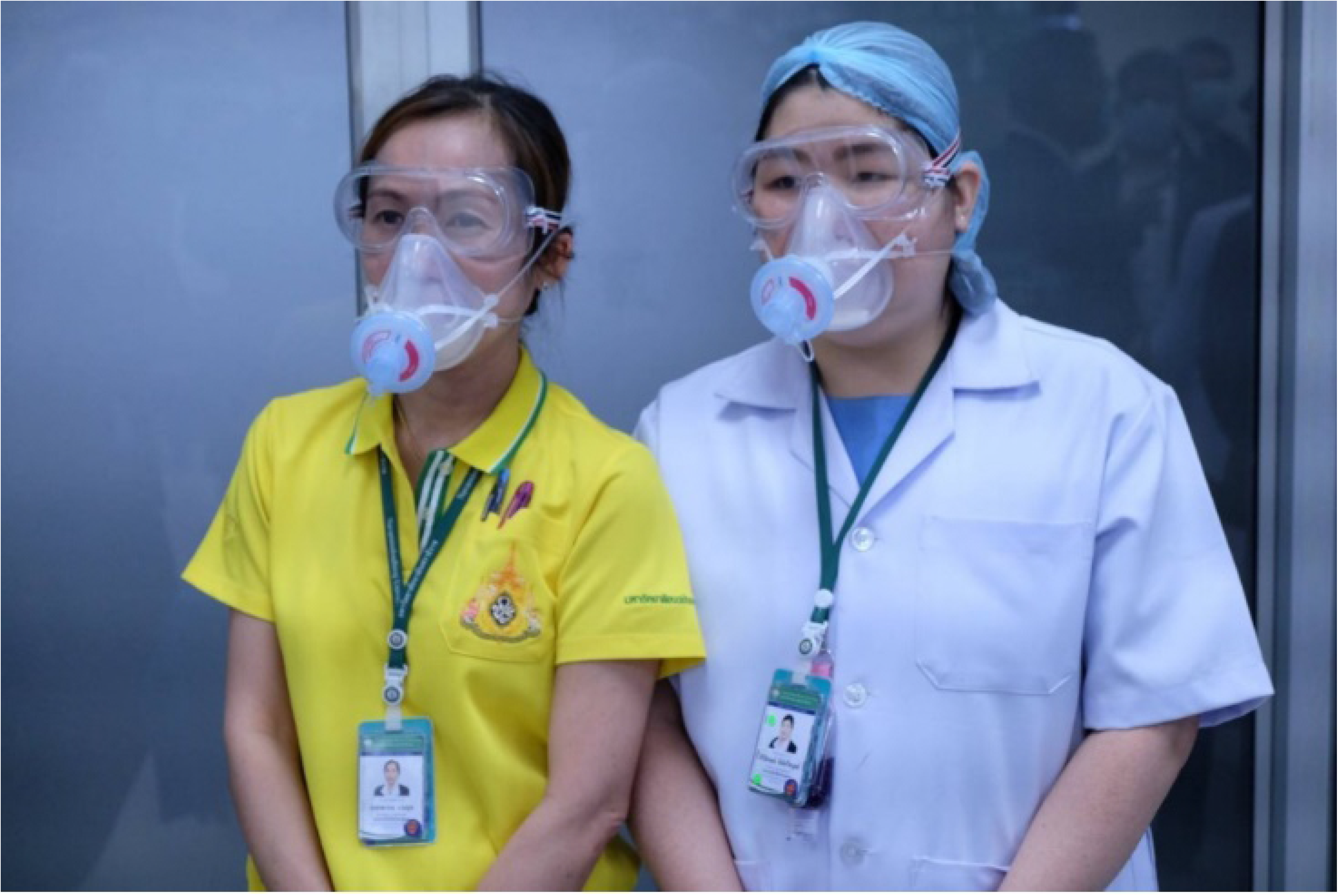
Health Care Workers Wearing Silicone N99 Masks.

As required by Occupational Safety and Health Administration OSHA) [7], the half piece respirators need to pass the fit test to identify those individuals who do not achieve a sufficiently good fit necessary for adequate protection. The performance of this respirator was also determined by measuring percent leakage under constant airflow [8]. The purposes of this study were to evaluate the fitting characteristic of our newly invented silicone N99 respirator and the performance against 30-μm particles using NaCl aerosols. The study was conducted to achieve two specific research objectives: (1) to investigate the fit characteristics of the novel silicone mask and whether the strap adjustment can help reduce the face-seal leakage; and (2) to determine the level of performance by measuring the inward leakage of the generated aerosols with a new filter and a used filter (for up to 24 h).

## Materials and methods

### Subjects

Participants were healthcare workers at Vajira Hospital, Bangkok, Thailand. The Vajira Institutional Review Board, Faculty of Medicine, Vajira Hospital, Navamindradhiraj University, approved the protocol, and all participants provided informed consent. The inclusion criteria were: healthy volunteers, 18 to 60 years old. The major exclusion criteria were: contraindications to fit tests, such as asthma, congestive heart failure, anosmia, and ageusia. Forty-three people (22 males, 21 females; mean age 28.71 ± 7.04 years) participated in this study. We excluded one male due to test intolerance (difficulty in breathing while putting on the hood cover). The other female was excluded because of a failed sensitivity test (no tests after 30 squeezes of sensitivity test solution). The remaining 41 people (21 males and 20 females) chose the proper size of silicone respirators to perform the qualitative fit testing.

### Filtering facepieces

The newly invented silicone half-piece respirators were tested. The configuration and model have been described above. There are three available sizes: small, medium, and large. None of the models tested in this study had exhalation valves. The CareStar filter is an electrostatic filter, and SafeStar is a HEPA filter product that is high performance, retaining at least 99.99% of bacterial and virus, with high hydrophobicity and transparent housing for visual control.

### The fit test procedure

A fit test was done with qualitative (Bittrex Solution aerosol, Qualitative Fit Test). The protocol was conducted in accordance with the protocol from the OSHA respiratory protection standard [9], including the number, type, and duration of the exercise, and the seal checks in accordance with the manufacturer’s instructions [10].

### Bitrex fit test

The Bittrex test uses a person’s ability to test a bitter solution to determine whether a respirator fits properly [11]. Each subject was given a taste-threshold screening test prior to each fit test to ensure that he or she could taste. This process was done without the subject wearing a respirator. After passing the sensitivity test, the subject will proceed through the seven steps of the fit test, as follows: breathe normally; breathe deeply; head side to the side; head up and down; bent over the wrist, jogging; talking; breath normally.

Each step took 60 s, and the Bitrex solution was refilled into the hood every 30 s, with half the dosage of the amount of the previous test. We asked the participant if he/she could taste the solution during each step of the test. The test was considered pass or fail.

We chose the qualitative fit test became it is more widely used [12], simpler to use, easy to transport [13], faster to perform, and cheaper to set up than the qualitative fit test [14].

Whenever the test failed, we would adjust the strap behind the respirator to tighten the face-seal leakage and repeat the procedure again. We would repeat the test twice before considering the test a failure.

Finally, we took note of the collected data, such as gender, age, size of the respirators, number of sprays, threshold level, and test result (pass, fail).

### Respirator performance

The real-time respirator performance test method was developed using a MT-05U machine (SIBATA model, Saitama, Japan), which measures particle concentrations of 0.03–0.06 μm diameter using a particle generator laser beam scattering particle counter, that measures particles outside and inside the mask. The ratio between these two values is considered the percentage leak, as follows:

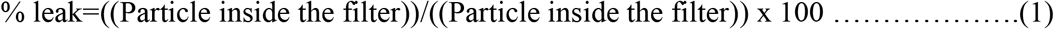

The percentage filter performance was calculated as 100 - % leak.

We tested at an airflow rate of 40 L/min, which is 3–4 times higher than normal physiology, assuming that there should be no leakage through the filter. We tested two types of filters, CareStar and SafeStar, that were incorporated into the silicone mask. CareStar is limited for 24-h usage, and SafeStar is for 72-h duration.

### Study protocol

We designed the filter test, as shown in Fig 4. We put the filter on the airflow generator, which generates an airflow of about 88 L/min. The NaCl aerosol condensation was generated by a MT-05 machine and flown through the cannula with a concentration of at least 70 particles/cc (the minimum level needed to conduct the test, in this study the average was 235 particles/cc) into the test chamber (green tube). We then measured the real-time percentage leak by counting the particles inside the filter compared to outside.

**Fig 4.**
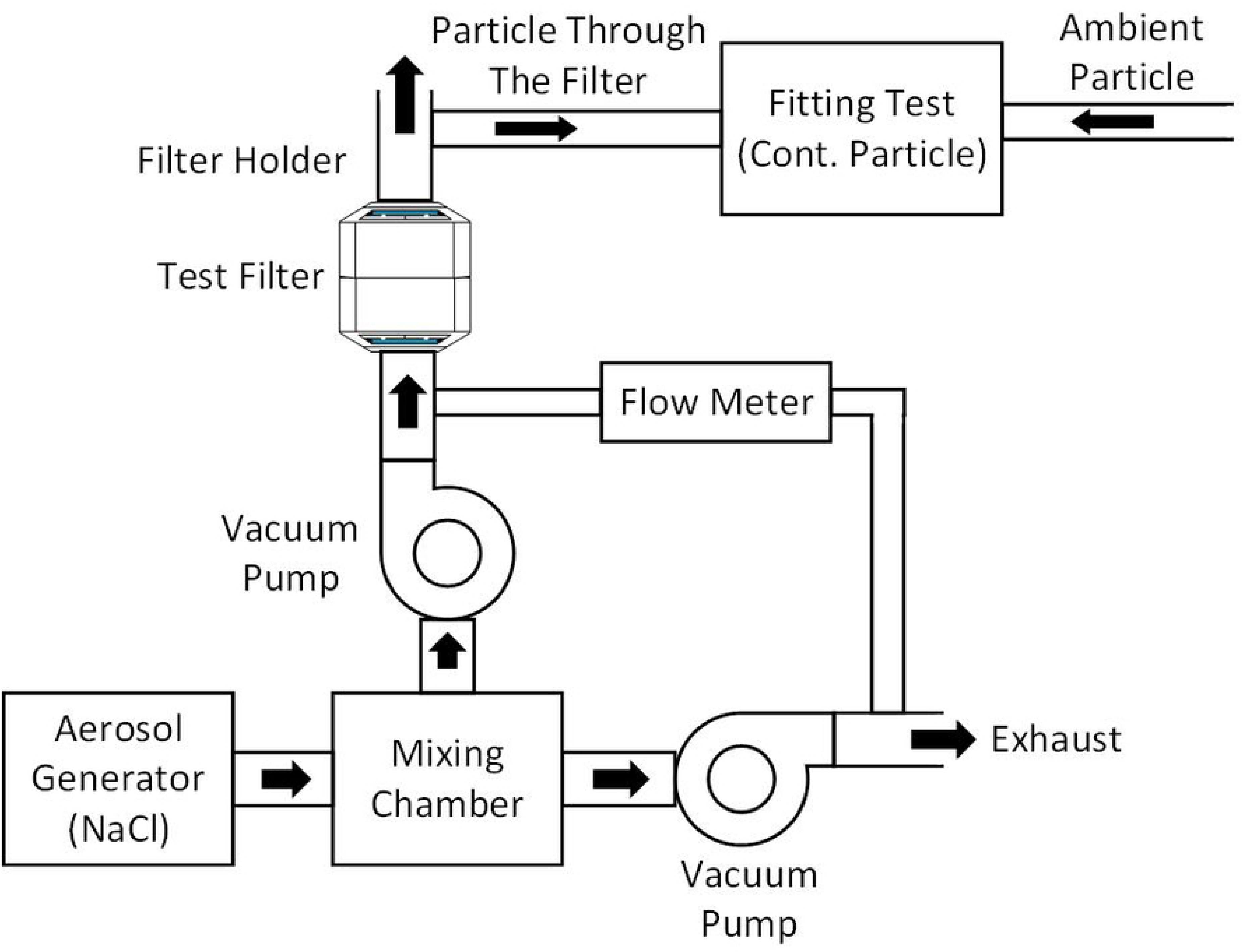
Diagram of the Filtration Test.

### Data collection

#### Sample size calculation

##### Fit test

We proposed the assumption that all subjects should pass the fit test by the non-inferiority hypothesis. The sample size estimation can be done by comparing the ratio of one group to the constant number (tooling for one population proportion) for non-inferiority trial as follows:

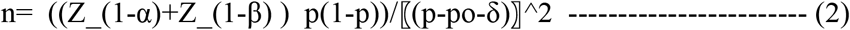

where n is the sample size, Z_(1 − α⁄2) is the statistical number under normal distribution according to ∝ = 0.05, and thus Z_(1−α) = 1.645, p is the ratio of sample size that pass the test at 99%, po is the reference value (100% Pan the test), and δ is the statistically significant value at 0.05.

Therefore:

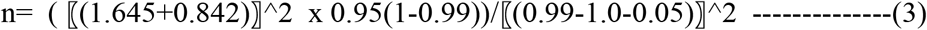

n=39

We included at least 39 persons.

##### Filter performance

We calculated the sample size using the test for non-inferiority for testing two dependent means as follows:

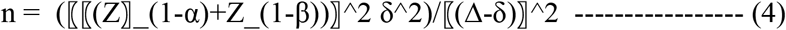

where Δ is the difference of mean of the population, δ is the statistically significant value, and δ^2 is the variance.

Since there are no previous studies to cite for references, we have used the G power version 3.1.9.4 program to compute filter performance. Two groups were compared using a paired t-test, where ∝ = 0.05 (one tall) and the power of the test is 80%. We applied effect size to 0.5 (medium effect size) [15].

n=27

We used the panel N99 respirators, which were used from 1 to 24 h to test for the filtration performance, relative to a control (new filter) respirator.

### Data analysis

The data analysis was performed using SPSS version 22.0 and Microsoft Office Excel for presenting the demographic data and respirator size. Descriptive statistics were calculated. The normal distribution of the data was tested using on the Shapiro-Wilk Test. The fittest passing rates (i.e., the number of subjects passing each fit test divided by the total number of subjects performing that fit test) were calculated. Filter penetration data were reported and calculated as mean ± SD, and compared to a specified target protection value of 99% in our respirator model.

## Results

Table 1 provides a summary of the baseline demographic of all participants. Forty-three subjects entered the study (22 male, 21 female). We excluded two persons due to test intolerance (fatigue during the test) and insensitive taste. The remaining 41 subjects (21 male, 20 female), mean age 28.22 ± 6.34 years old, used three sizes of N99 respirators: small (n = 10), medium (n = 25), large (n = 6) (Table 2).

**Table 1.** Baseline Characteristics of Participants Performing Fit Test (n = 41).

| Characteristics |  |
| --- | --- |
| Gender, n (%) |  |
| Male | 21 (51.2) |
| Female | 20 (48.8) |
| Age (year), mean±SD | $28.22 \pm 6.34$ |
| BW (kg), mean±SD | $64.87 \pm 20.10$ |
| BMI (kg/m<sup>2</sup>), mean±SD | $23.20 \pm 5.49$ |
| Mask size |  |
| S | 10 (24.4) |
| M | 25 (61.0) |
| L | 6 (14.6) |
| Number of tests, median (IQR) | 10 (5–12) |
| (min - max) | (3–30) |

**Table 2.**
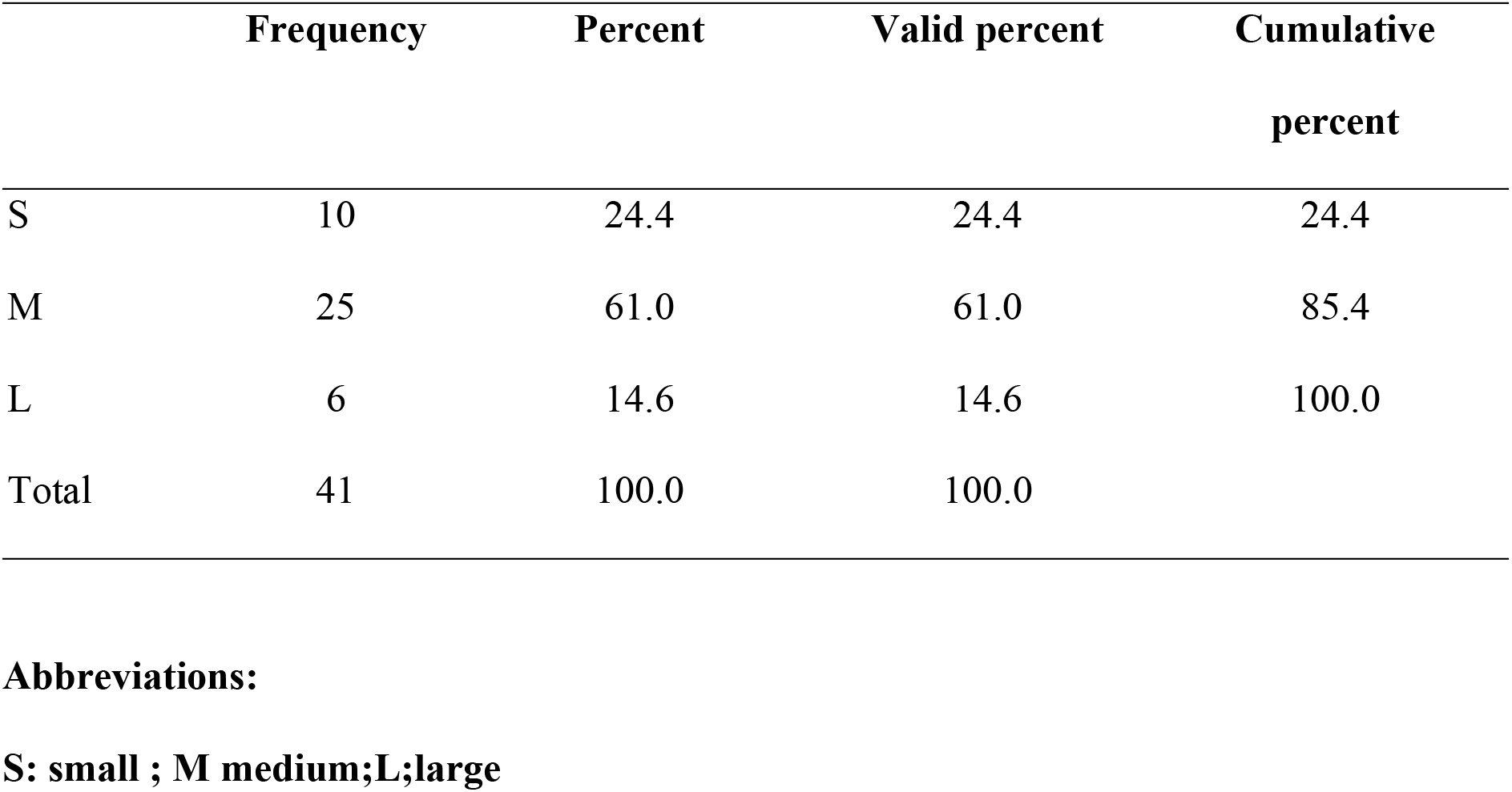
Distribution of Mask Sizes.

Thirty-two subjects (78%) passed the first fit test. After adjusting the O-ring trap to tighten the respirators, seven subjects passed the second test (80%). There was one person who did not pass the test, even after adjusting the strap for the third time. The overall fit-test passing rate in this study was 40/41 (97.6%) (Tables 3 and 4).

**Table 3.** Summary of the Fit Test Results (N=41).

| Fit test | Result |  |  |  |
| --- | --- | --- | --- | --- |
|  | Pass |  | Failed |  |
|  | n | Percent | n | Percent |
| 1<sup>st</sup> Fit (n=41) | 32 | 78.0 | 9 | 22.0 |
| 2<sup>nd</sup> Fit (n=9) | 7 | 77.8 | 2 | 22.2 |
| 3<sup>rd</sup> Fit (n=5) | 4 | 80.0 | 1 | 20.0 |
| Cumulative result (n=41) | 40 | 97.6 | 1 | 2.4 |

**Table 4.** Data of Percent Filtration of Each Filter.

| No. | Type | Outside | Inside | % Filter<br>performance | Time of use<br>(hr) | Amount | % Leak |
| --- | --- | --- | --- | --- | --- | --- | --- |
| 1 | Safe Star | 7689 | 2 | 99.97 | 1 | 1 | 0.026 |
| 2 | Safe Star | 8374 | 2 | 99.98 | 1.3 | 1 | 0.023 |
| 3 | Safe Star | 8633 | 2 | 99.98 | 2 | 1 | 0.023 |
| 4 | Safe Star | 7681 | 2 | 99.97 | 2 | 1 | 0.026 |
| 5 | Safe Star | 8048 | 4 | 99.95 | 3 | 1 | 0.049 |
| 6 | Safe Star | 12871 | 17 | 99.87 | 3 | 1 | 0.132 |
| 7 | Safe Star | 7746 | 0 | 100 | 4 | 1 | 0 |
| 8 | Safe Star | 14718 | 4 | 99.97 | 5 | 1 | 0.027 |
| 9 | Safe Star | 5553 | 11 | 99.8 | 8 | 1 | 0.198 |
| 10 | Safe Star | 11676 | 4 | 99.97 | 8 | 1 | 0.034 |
| 11 | Safe Star | 7646 | 0 | 100 | 9 | 1 | 0.09 |
| 12 | Safe Star | 8523 | 3 | 99.96 | 10 | 1 | 0.035 |
| 13 | Safe Star | 12105 | 0 | 100 | 10 | 1 | 0 |
| 14 | Safe Star | 12422 | 14 | 99.89 | 24 | 1 | 0.112 |
| 15 | CareStar | 7986 | 129 | 98.38 | 2 | 1 | 1.615 |
| 16 | CareStar | 12092 | 190 | 98.43 | 4 | 1 | 1.57 |
| 17 | CareStar | 9199 | 101 | 98.9 | 10 | 1 | 1.09 |
| 18 | CareStar | 12754 | 200 | 98.4 | 10 | 1 | 1.56 |
| 19 | CareStar | 12356 | 14 | 97.64 | 24 | 1 | 0.113 |
| 20 | CareStar | 10651 | 213 | 97.25 | 1 | 1 | 0.08 |
| 21 | Safe Star | 8345 | 0 | 100 | 3 | 1 | 0 |
| 22 | CareStar | 2577 | 53 | 97.94 | 1 | 1 | 2.06 |
| 23 | Safe Star | 2694 | 0 | 100 | 30 | 1 | 0 |

### Filter penetration

Filler penetration for monodisperse aerosols in the 0.03-micron range aerosols was measured at a 40 L/min constant flow rate using the SIBATA MT 05U Fit test under the fit check mode (standard OSHA 29 CFR 1910.134). Table 4 shows the filler penetration of two filler types (SafeStar and CareStar, Draeger) [9]. We tested the baseline filter performance (first use) and used the filters for between 1 and 24 h. All of the filters had at least 99% protection (mean, 99.96 ± 0.06 for SafeStar, and 98.13 ± 0.56 for CareStar, p < 0.001). The SafeStar filter had better efficiency than CareStar (Table 5). Even with a use time of up to 24 h, the protection value remained > 99% (Fig 5).

**Table 5.** Mean Filtration Efficiency of the CareStar and SafeStar Filters.

| Variables | Total | SafeStar | CareStar | P-value |
| --- | --- | --- | --- | --- |
|  | 23 (100) | 16 (69.57) | 7 (30.43) |  |
| Particle outside, Mean±SD | 9232.13±3129.9 | 9045.25±3019.5 | 9659.29±3580.5 | |
|  | 0 | 0 | 4 |  |
| (Min-Max) | (2577-14718) | (2694-14718) | (2577-12754) |  |
| Inside, Mean±SD | 41.96±71.19 | 4.06±5.26 | 128.57±77.03 | |
| (Min-Max) | (0-213) | (0-17) | (14-213) |  |
| % Performance Filter, Mean±SD | 99.40±0.91 | 99.96±0.06 | 98.13±0.56 | <0.001 |
| (Min-Max) | (97.25-100) | (99.80-100) | (97.25-98.90) |  |

**Fig 5.**
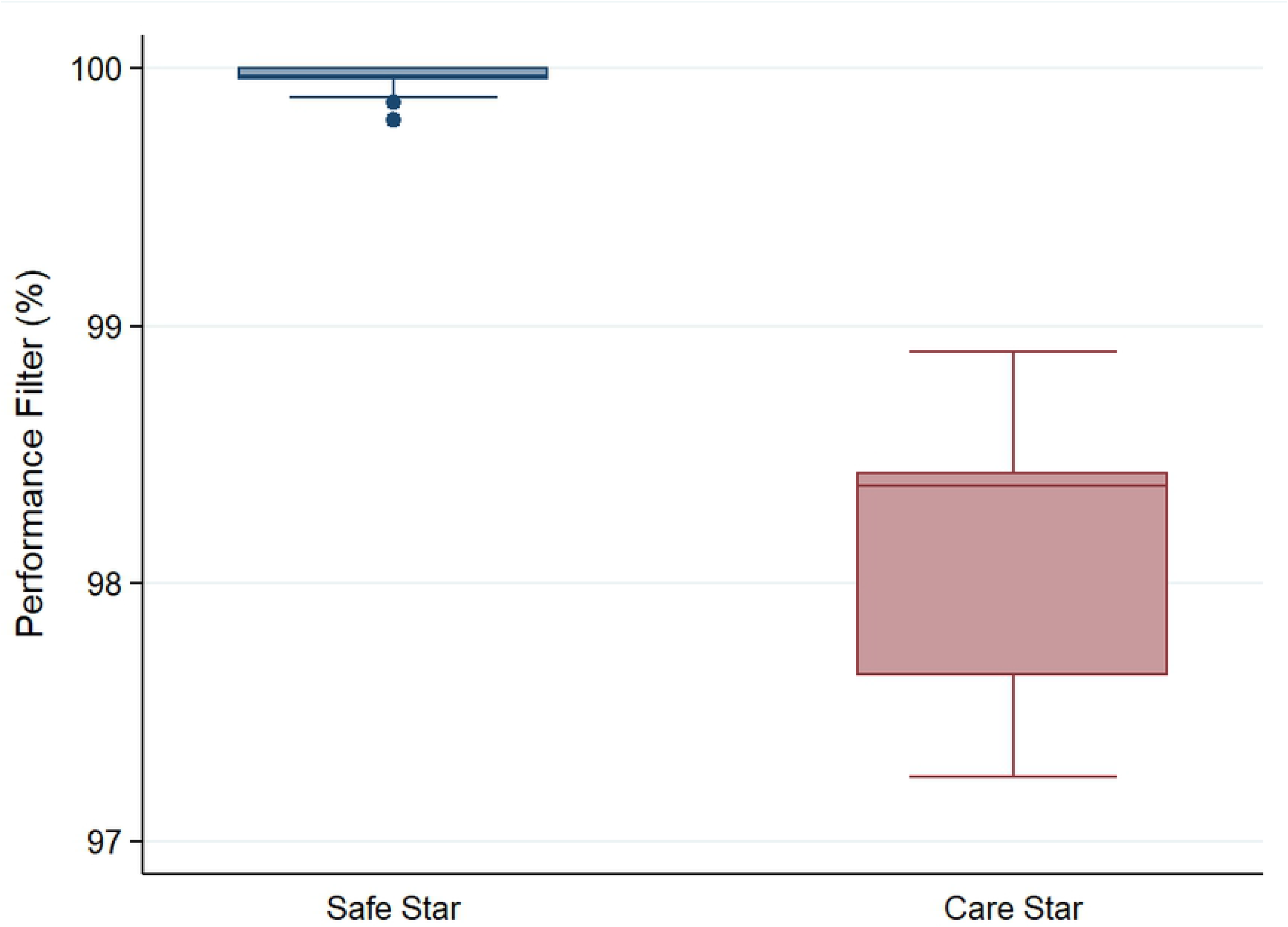
Comparison between the Percentage Performances of the SafeStar and CareStar Filters.

## Discussion

It is widely accepted that wearing face masks in public corresponds can help to prevent inter-human transmission of SARs-CoV2 [16]. In HCP that remain at risk to COVID-19, patients should follow appropriate infection control procedures. These levels of protection depend on the setting where modes of viral transmission are relevant. Filtering facepiece respirators are essential devices to protect HCW from bioaerosol particles [17,18]. According to the National Institute for Occupational Safety and Health (NIOSH) regulations 42 CFR 84, the N 95 respirators are recommended for personal protection from exposure to respiratory aerosol particles [19]. However, due to the COVID-19 pandemic, there is a global supply shortage for the most exposed persons, HCWs. This underlines the urgent need to replenish the masks using a variety of available materials.

Our team has invented a novel N99 half-piece respirator by combining a silicone facemask (as used in the operating rooms) with an electrostatic filter plug (CareStar or safe star Draeger Lubeck, Germany). We developed an O-ring strap made from silicone, similar to the silicone mask, to tighten the mask to the face. The strap consists of three parts: the silicone strap; the strap locker to adjust the length (made from polypropylene); and a 4-way hook (made form silicone to connect the strap with the mask).

By adjusting the O-ring straps, almost all patients passed the fit test. This shows a good level of protection. Adjustable head straps then allow a better-customized seal because they can be tightened to better secure the respirators. At first, three different sizes of the face pieces were produced. The majority of our participants used medium sizes. It should be noted that fit-testing is just one factor to determine the level of protection provided by a respirator. Other tests, such as filter penetration, should also be done. According to the specification of the electrostatic filter (CareStar or SafeStar), the estimated filtration efficiency should be more than 99% after testing with the SIBATA machine. The new filter had the filtration efficiency of more than 99%. We then evaluated the filters after use (1 to 24 h) since the filter efficacy might be hampered by humidity and the formation of a biofilm layer over the filter surface. The test protocol that we adapted measures the percent leakage through the filters by counting the sodium chloride generated aerosols. The percent of leakage was less than 1%. Thus, we can confirm that the efficiency of our new respirators was compatible with the N99 type of respirator and can be used at least up to 24 h before changing the filter.

## Limitations

The limitations of this study were the method of fit test, that was qualitative and should be confirmed with qualitative fit test7,8. The silicone mask still has some drawbacks, such as the hardness of the material, that is comfortable for the wearer. Those who wear eyeglasses might have difficult place his/her eyeglasses on the bridge of the silicone mask. The levels of protection afforded by the respiratory protective devices in this study might not be representative of all respirator wearers, who might have different facial size distributions than those of the subjects in this study. The filters that we tested had limited use times (up to 24 h), while their efficiency remained excellent. Further tests should be performed for a filter that has been used for more than 24 h. The filtration test was designed with the equipment available in the unit based on the urgency to search for alternative protective devices.

Despite the limitations of our study, a strength of this innovation is that the materials are available, and the efficacy is acceptable, and even superior to the long-used N95 masks.

## Conclusions

Here we report the efficacy of our innovation, a silicone N99 mask that we have named VJR-NMU N99 half-piece respirators. The VJR-NMU N99 respirators surpassed the expected levels of protection, and can be useful in the context of a global shortage of PPE. However, this is the first version of our masks, and further modifications are needed to improve user friendliness and provide adequate protection. We believe that the findings of this study will contribute to the provision of safe and superior healthcare services for HCW, and that the VJR-NMU N99 respirators can help to replenish the shortage of essential healthcare worker protection.

